# ATLIGATOR: Editing protein interactions with an atlas-based approach

**DOI:** 10.1101/2022.01.19.476980

**Authors:** Josef Paul Kynast, Felix Schwägerl, Birte Höcker

## Abstract

**Motivation:** Recognition of specific molecules by proteins is a fundamental cellular mechanism and relevant for many applications. Being able to modify binding is a key interest and can be achieved by repurposing established interaction motifs. We were specifically interested in a methodology for the design of peptide binding modules. By leveraging interaction data from known protein structures, we plan to accelerate the design of novel protein or peptide binders.

**Results:** We developed ATLIGATOR – a computational method to support the analysis and design of a protein’s interaction with a single side chain. Our program enables the building of interaction atlases based on structures from the PDB. From these atlases pocket definitions are extracted that can be searched for frequent interactions. These searches can reveal similarities in unrelated proteins as we show here for one example. Such frequent interactions can then be grafted onto a new protein scaffold as a starting point of the design process. The ATLIGATOR tool is made accessible through a python API as well as a CLI with python scripts.

**Availability and Implementation:** Source code can be downloaded at github (https://www.github.com/Hoecker-Lab/atligator), installed from PyPI (“atligator”) and is implemented in Python 3.

## 1 Introduction

For protein design it is crucial to understand how proteins form interactions. Interactions can be formed intramolecularly to define stability or function as well as intermolecularly with various interaction partners such as solvent, small molecules, peptides, or full proteins. Thus, the choice of a particular amino acid at a certain position in a protein is crucial to establish such favorable interactions between two or more amino acid residues. Hence, understanding how specific residue types interact with each other is of particular interest when creating newly designed proteins.

A description of the conformational space that is occupied by interacting amino acid side chains in known protein structures as well as relative positioning of both interaction partners can provide powerful information for protein design. Singh and Thornton (Singh and Thornton, 1992) already classified interactions between pairs of distinct residue types. Moreover, they described clusters of orientation and position combinations within these pairs of amino acids. In a similar approach, Vondrasek and colleagues investigated interaction energies of amino acid combinations calculated in gas-phase (Berka *et al*., 2009, 2010; Galgonek *et al*., 2017). For the analysis of enzymatic active sites, groups of amino acid residues from three-dimensional structures were categorized, based on sequence alignments (Porter *et al*., 2004). While these studies led to a better understanding of amino acid interactions, they were constrained by the available input data at that time as well as by a focus on analysis rather than application in the design of binding features.

Despite this idea being around for three decades, favored interactions of amino acids are currently not used intensely in rational protein design approaches. However, the usage of interaction modes that can be found frequently in nature and thus appear more favorable than others might help to shape the interaction of polypeptides. Such information allows to modify not only internal interactions within a protein, but also interactions with a different peptide or protein binding partner. Furthermore, the identification of frequently interacting residue groups plus their favored conformations opens the possibility to graft specialized binding pockets to specifically bind peptide or protein targets of interest.

During the development of this method for the design of protein-peptide interactions we had a particular application in mind, namely the creation of custom-made peptide-binders based on armadillo repeat proteins stitched together from repeat modules. Armadillo repeat proteins comprise a natural binding interface for elongated peptide stretches which was further refined to exhibit peptide binding in a regular fashion (Hansen *et al*., 2018, 2016). Thus, grafting existing binding modes of known structures on the binding interface of such a single repeat would be a crucial step to design new modules that can be assembled or incorporated in an existing peptide binder.

To enable such rational design, we now present the software tool ATLIGATOR, short for ATlas-based LIGAnd binding site ediTOR. It allows the user to analyze frequent interaction modes of two or more amino acids and to directly apply this information to rational design approaches. The program relies on data structures called *atlases* that contain descriptions of pairwise interactions from protein structures. A collection of structures that builds up such an *atlas* is a subgroup of all structures in the Protein Data Bank (Berman *et al*., 2000) and can for example represent a certain type of fold based on classifiers of the SCOPe database (Fox *et al*., 2014). Moreover, the ATLIGATOR tool also incorporates association rule learning in the form of frequent itemset mining to extract frequent groups of pairwise interactions based on single ligand residues from the *atlas*. These groups are called *pockets* and represent starting points for protein interface design tasks. This representation is based on the assumption that favorable interaction groups have been established during the evolution of the proteins of choice and are thus detected as *pockets*. A major key functionality of ATLIGATOR is the ability to visualize each individual step of the ATLIGATOR toolchain interactively. Furthermore, ATLIGATOR *atlas* and *pocket* datapoints can not only be browsed for individual amino acid combinations but can additionally be used in an integrated tool called Manual Design. Manual Design allows to use a protein-peptide complex structure of choice and alter the interaction surface by binding pocket grafting or manual mutations with recommendations based on *pocket* data. Hence, ATLIGATOR acts as a framework that offers a multitude of possible workflows. Besides the use of single parts for analysis or design, the setup offers a complete workflow from the analysis of interaction modes in protein structures all the way to the interactive application of protein interface design by leveraging previously accumulated knowledge.

## 2 System and Methods

### 2.1 Algorithms

ATLIGATOR is a versatile toolkit for the analysis and the design of protein interactions. It focusses on single side chains of one interacting partner (the ligand) and its relation with multiple residues at the surface of the other interacting partner (the binder) that form a binding pocket for the single residue. *Atlases* are generated for all of these interactions within a user-specified set of complex structures. Through this focus on the single residue interaction level, the tool allows to detect promising interaction features. This knowledge can directly be applied to specific design problems of protein complex interfaces. The toolkit contains the following parts.

#### 2.1.1 Structure selection and preprocessing

The information gathered by ATLIGATOR is extracted from existing protein structures derived from the Protein Data Bank (PDB). The PDB contains an abundant collection of protein structures, which have been derived mainly from experimental methods. It is useful to be able to select the qualitatively best structures as well as the most fitting structures, e.g. from the same protein family, fold or class. Therefore, we provide the option to select structures based on one’s own rationale or on identifiers of the SCOPe database, thereby creating sets of structures with shared structural or evolutionary background.

Furthermore, we allow to additionally filter structures for certain properties and quality criteria using a pre-selection and processing utility. This utility is called ALARMS, short for Automated Ligand ARMadillo protein pre-Selector, and is capable of applying the following filter criteria:

1. Specific protein families (e.g., by SCOPe query)
2. Minimum/maximum length of binder and ligand sequences
3. Maximum distance between ligand and any binder residue
4. Secondary structure content

ALARMS produces a directory of pre-processed pdb files, each containing one ligand-binding complex where ligand and binder are located in individual chains, removing unnecessary parts of ligand chains. These files are then used for *atlas* generation after an optional filtering step.

#### 2.1.2 Atlas generation

The pre-selected input structures contain external coordinates for the atoms of different ligand residues and binder proteins, respectively. An *atlas* is a collection of filtered and transformed datapoints, each describing an interaction between one residue of the ligand and one residue of a binder. The following algorithm describes on a coarse level how *atlas* datapoints are obtained from the input structures:

##### Algorithm 1

Atlas Datapoint Extraction as Simplified Python Code.

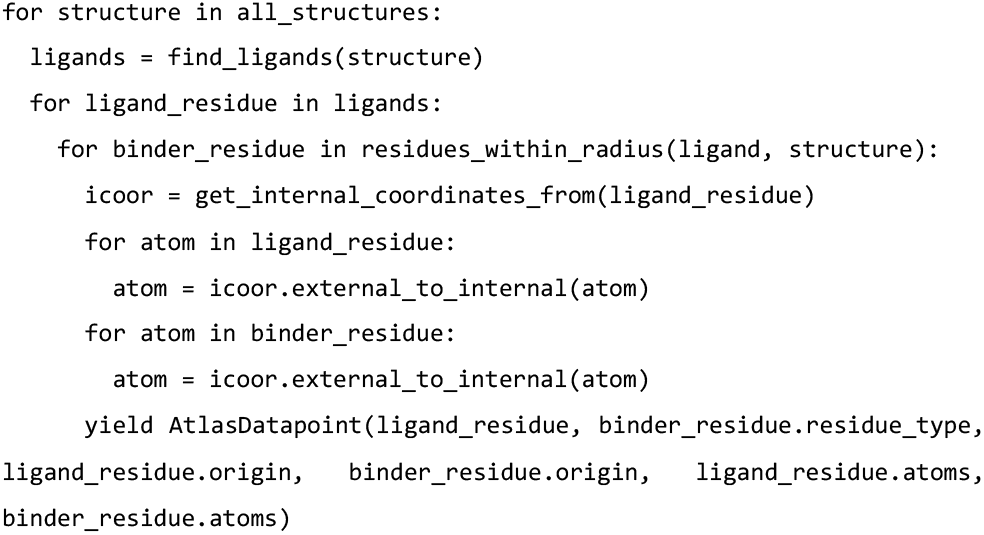

For determining whether any pair of ligand and binder atoms are considered as interacting, we define specific interaction distances.

These distances depend on the type of interactions between ligand and binder atom:

- Ionic: interactions between positively and negatively charged atoms (default: 8.0 Å).
- Aromatic: interactions between carbon atoms of aromatic rings (default: 6.0 Å).
- Hydrogen bonds: interactions between donor and acceptor atoms (default: 6.0 Å).
- All other interactions, e.g., hydrophobic (default: 4.0 Å).

The interacting residues are transformed into an internal coordinate system, which allows to detect patterns in pairwise interactions, seen from the perspective of ligand residues. It is defined as follows:

- The ligand residue’s *C*_α_ atom is the origin.
- The ligand residue’s *C*_β_ atom is located on the x-axis of the internal coordinate system.

(For glycine, we simulate a virtual *C*_β_ atom for this purpose.)

- The ligand residue’s C atom (carbonyl carbon) lies within the xy-plane.
- The ligand residue’s N atom is defined with a negative z-value.

Every *atlas* is composed of datapoints storing individual interactions between two residues – a ligand and a binder residue. This collection of datapoints is grouped into *atlas pages* including all datapoints of a certain ligand residue type. *Atlas pages* are partitioned further into *atlas maps* including all datapoints of a combination of one ligand residue amino acid type interacting with one binder residue amino acid type (Figure 2).

#### 2.1.3 Spatial similarity function

To compare *atlas* datapoints with each other or with designable binder residues we created a distance-orientation function to describe the spatial similarity of two residues *R*_1_ and *R*_2_. Assuming that they are both represented in the same, internal or external, coordinate system, their distance |*R*_1_ − *R*_2_| is defined as follows:

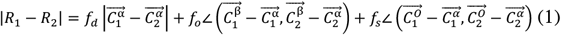

The equation considers positions of Cα atoms of both interacting residues (where 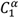 denotes the *C*_α_ atom of *R*_1_, etc.). Furthermore, the angles between two characteristic orientation vectors, namely those between *C*_α_ and *C*_β_ (referred to as primary orientation below) as well as *C*_α_ and the carbonyl C (secondary orientation) of the residues are compared. The weight factors *f*_d_, *f*_0_ and *f*_s_ can be adjusted by the user; the default values are 1.0Å^− 1^ for *f*_d_ and 2.0 for both *f*_0_ and *f*_s_.

#### 2.1.4 Pocket mining

Ligand-binder interactions as shown in the *atlas* do not have a purely pairwise nature. Several binder residues can instead contribute to binding one ligand residue. If similar binder residue groups form interactions to ligand residues in various structures, interaction patterns can be extracted and generalized. We call such a frequently occurring interaction pattern a *pocket*. Such *pockets* can be detected and extracted from an *atlas* database which is described below.

##### Itemset Extraction

In its first step, the algorithm exploits the fact that datapoints of the *atlas* include their origin. Hence, we group all datapoints originating from the same ligand residue. To detect which *pockets* are frequent, we reduce the information stored in these groups into natural itemsets, which are mere enumerations of binder residues that interact with the same ligand residue.

##### Frequent Itemset Mining

Depending on the size of the *atlas*, we obtain a large number of itemsets for every specific ligand residue type in this way. In the field of business intelligence, the so-called a-priori algorithm (Agrawal *et al*., 1993) has been established; we apply this procedure in order to find representative subsets that are contained in a relevant share of natural itemsets extracted from the *atlas*. As a result, these subsets are groups of binder residue types that are found to interact frequently with a ligand residue type.

##### Pocket Extraction

Frequent itemsets indicate which residues are part of a *pocket*, but ignore their structure. This information in turn is added during *pocket* extraction where the coordinates of the underlying *atlas* datapoints play a major role. In this step, natural pockets of the *atlas* are matched with frequent itemsets in order to identify those *pockets* that represent the itemset (i.e., they include the same collection of residues or more). This adds the structural component of each pairwise interaction from the *atlas* to the group of amino acid types within the itemset. The result of this analysis is stored in a superimposed way; technically, every superimposed *pocket* is a subset of the original *atlas*.

##### Clustering, Noise Reduction, and Selection of Representative

Last, the information stored in every superimposed *pocket* is clustered. To this end, we employ a modified variant of the k-means algorithm (Chen *et al*., 2004), utilizing the spatial similarity function shown in Section 2.1.3 for the calculation of cluster centroids and variances (i.e., the mean deviation of clustered *pocket* residues from the cluster centroid).

In order to reduce noise, the specific algorithm utilized here additionally ensures that the variance of a cluster is kept below a user-defined threshold (default value: 5.0 Å). By removing the most distant members from the cluster until the threshold is met, the noise present in superimposed *pockets* is reduced. For every cluster, we ultimately select as the most representative element (also called medioid) the natural pocket with the least distance from the cluster centroid. This is an instance of the so-called assignment problem, which can be solved using the Hungarian Algorithm (Kuhn, 1955).

#### 2.1.5 Pocket grafting

*Pocket* grafting is a simple method that directly exploits the information available from the *pockets* for the creation of designed ligand-binder complexes based on a scaffold. It takes the best-matching *pocket* residues of a selected *pocket* according to the spatial similarity function (see Section 2.1.3) and applies corresponding mutations to binder residues. The details of the procedure are described by the following algorithm:

##### Algorithm 2

Pocket Grafting as Simplified Python Code.

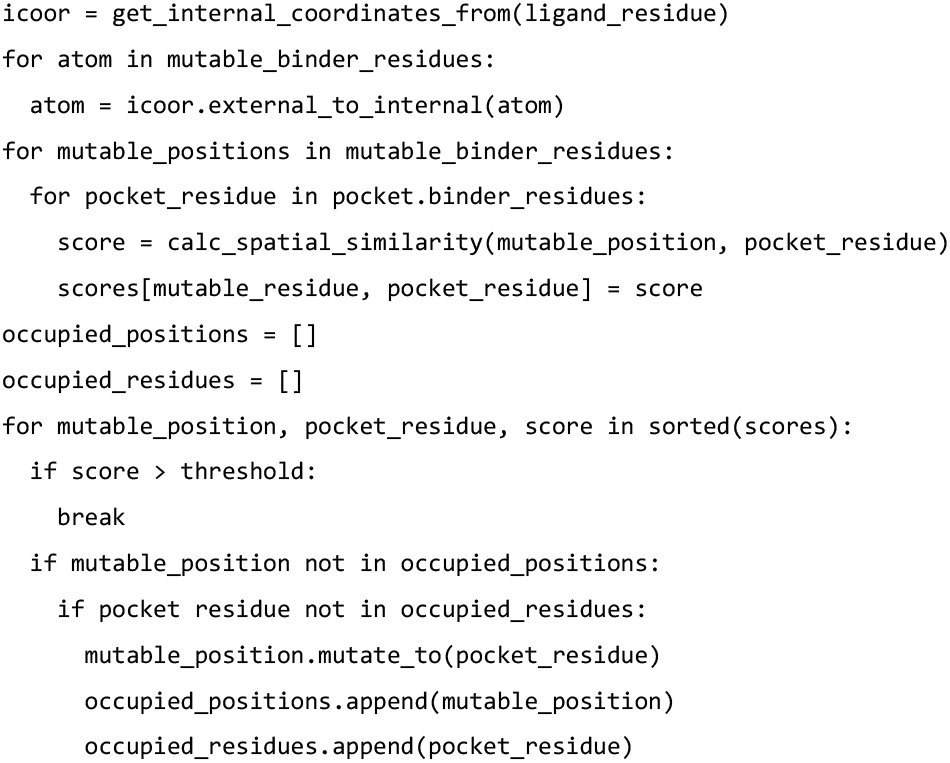

The algorithm contains an additional adjustable parameter, the distance threshold θd (default: 12.0), which prevents the alignment of bad-matching *pocket* residues.

#### 2.1.6 Quick graft

Pocket mining usually results in several *pockets* for each ligand residue type. To overcome the need to graft each *pocket* onto the scaffold individually and to select the best graft manually the quick graft protocol includes automatic grafting of the best matching pocket. To select the best pocket quick graft picks the pocket graft resulting in the best cumulative spatial similarity (see 2.1.3).

In the process of redesigning an interaction interface more than one ligand residue might be mutated. In this case, the grafted binding pockets need to complement each other to create the best fit for all exchanged ligand side chains. As a solution, quick graft detects conflicting grafts and finds the optimal set of pocket grafts with mutually exclusive positions to mutate. Additionally, the best n grafted designs can be generated to give the user the option to compare and select the best grafts.

### 2.2 Implementation

The algorithms discussed in Section 2.1 are implemented in Python 3. To access the functionality, we deliver scripts with user argument parsing as well as a raw python API (see Figure 1).

**Figure 1:**
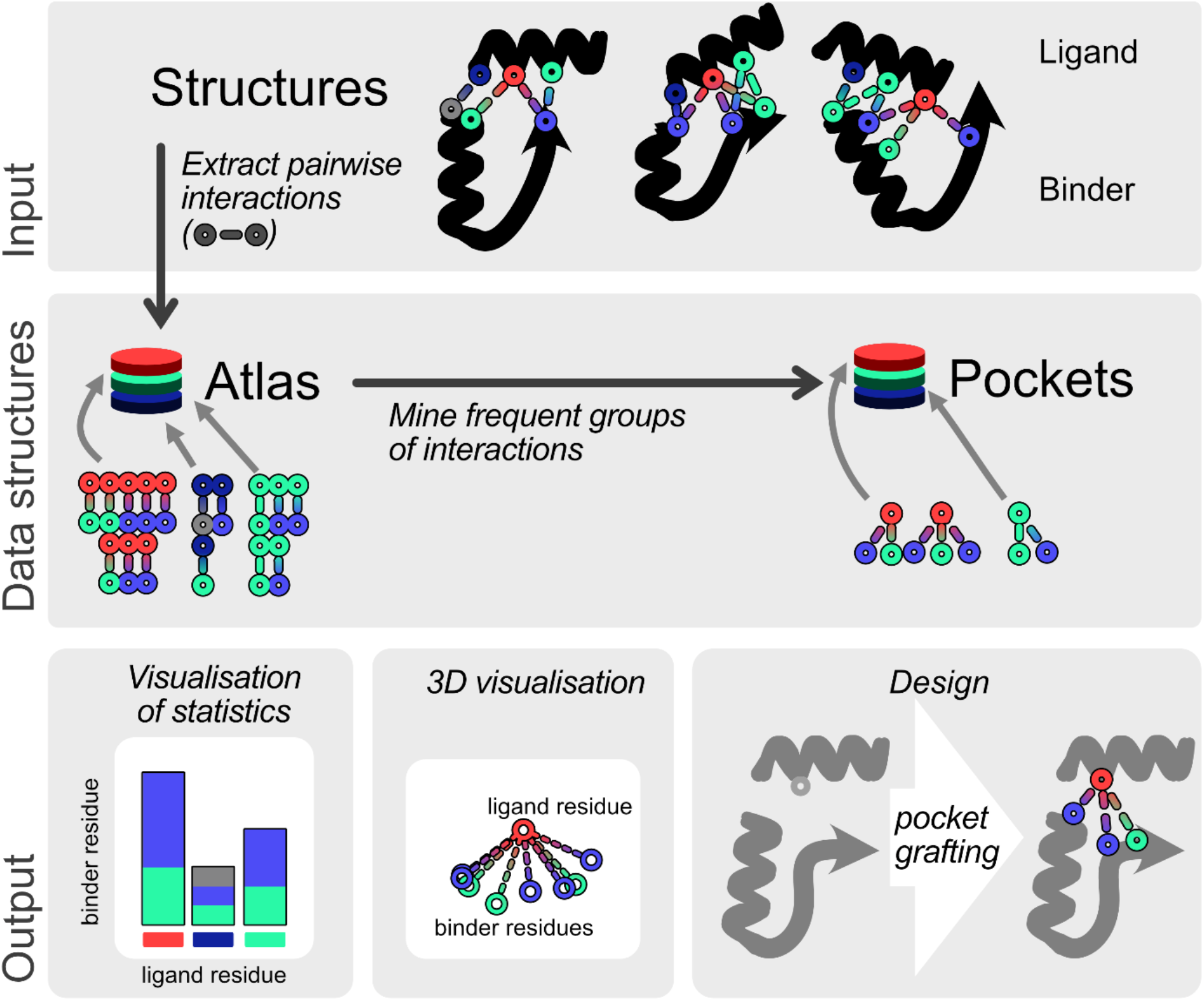
Figure 1: Overview of ATLIGATOR toolchain. The python-based tools of ATLIGATOR include the extraction of pairwise interactions from a *structure collection* as well as mining of frequent groups of interactions. Those tools as well as the input and output data can be accessed via a python API, meaning the source code as well as predefined scripts. Both types of interfaces can be used to analyze extracted interactions to find patterns which can be employed for new designs. This can be achieved by visualizing *atlas* statistics or 3D plotting of *atlas* and *pockets*. Moreover, ATLIGATOR includes the option to design new interaction sites based on binding pocket grafting.

#### 2.2.1 Python API and CLI

The API as well as the supplemented python CLI (command-line interface) allow to generate and access all parts of the ATLIGATOR toolchain, visualize *atlases, pockets* and additional statistics and follow preprocessing steps with ALARMS. Furthermore, within the API the visualization of single pockets and pocket grafting functions are available. The following paragraphs will guide through a typical workflow of the different tools.

##### Structure Selection

The PDB is a rich resource for protein structures. Due to the large amount of data, but also due to biases in structures, scanning the whole PDB for ligand-binder interaction information is not recommended. Rather, the user can select structures from the PDB based on own rationale or on identifiers of the SCOPe database, creating sets of structures with shared structural or evolutionary background. Furthermore, we allow to use preprocessing and filtering structure files to include ALARMS procedures (see 2.1.1). Those sets of structures are called *structure collections*.

##### Atlas Visualization and Usage

*Atlases* can be obtained with the *atlas* generation algorithm (see 2.1.2) from *structure collections. Atlases* do not only serve as input for further analysis and design but visualizing them directly also provides insights into the collected data. Of particular interest are the 20 different *atlas maps* encoded in every *atlas*, which show frequent interactions for given ligand residue types. ATLIGATOR offers a three-dimensional visualization of single ligand amino acid types against one or all other binder amino acid types, corresponding to *atlas maps* or *pages*. These plots contain Cα and Cβ atoms of the centered ligand residue as well as Cα and Cβ atoms of the binder residues of each included datapoint. Thus, information about the relative position as well as orientation of both interaction partners is provided. Furthermore, it provides statistical insights into the composition of the *atlas* in terms of pair-wise interactions such as frequency of detected interaction pairs.

##### Pocket Visualization and General Usage

*Pockets* can be mined directly from *atlases*. ATLIGATOR can visualize and export into pdb format both superimposed and representative *pockets* (see 2.1.4). To present a more detailed point of view *pockets* can also be plotted as a collection of all included datapoints, representing the *pocket atlas* as a filtered instance of the corresponding *atlas* page (see Figure 3A & B). Also, single *pocket* instances can be visualized, they contain exactly one ligand rotamer as well as all binder residue rotamers interacting with this exact ligand in the source structure as a part of this *pocket*. Thus, only those residues included in the *pocket* itemset that were not filtered during *pocket* generation will be present (see Figure 3C). *Pockets* constitute a useful information *per se*, but they are also utilized in an automated grafting algorithm.

**Figure 2:**
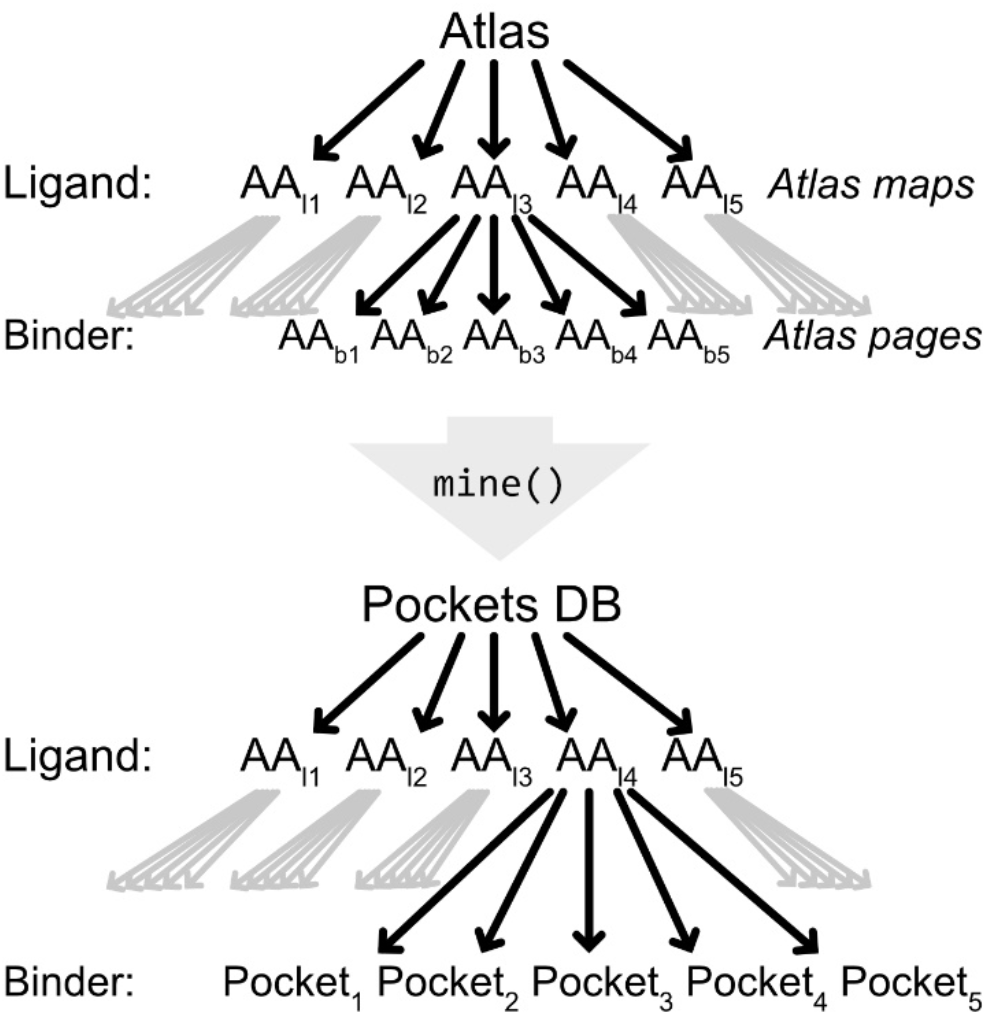
Structure of Atlas and Pocket Database. An *atlas* consists of *atlas maps* defining the ligand residue types that can be further subdivided into *atlas pages. Atlas pages* are defined by a specific ligand-binder residue type combination. *Pockets* are subgroups of the underlying *atlas* and can be structured by their ligand residue type as well. One *pocket* is defined by the ligand residue type in combination with a group of binder residue types.

**Figure 3:**
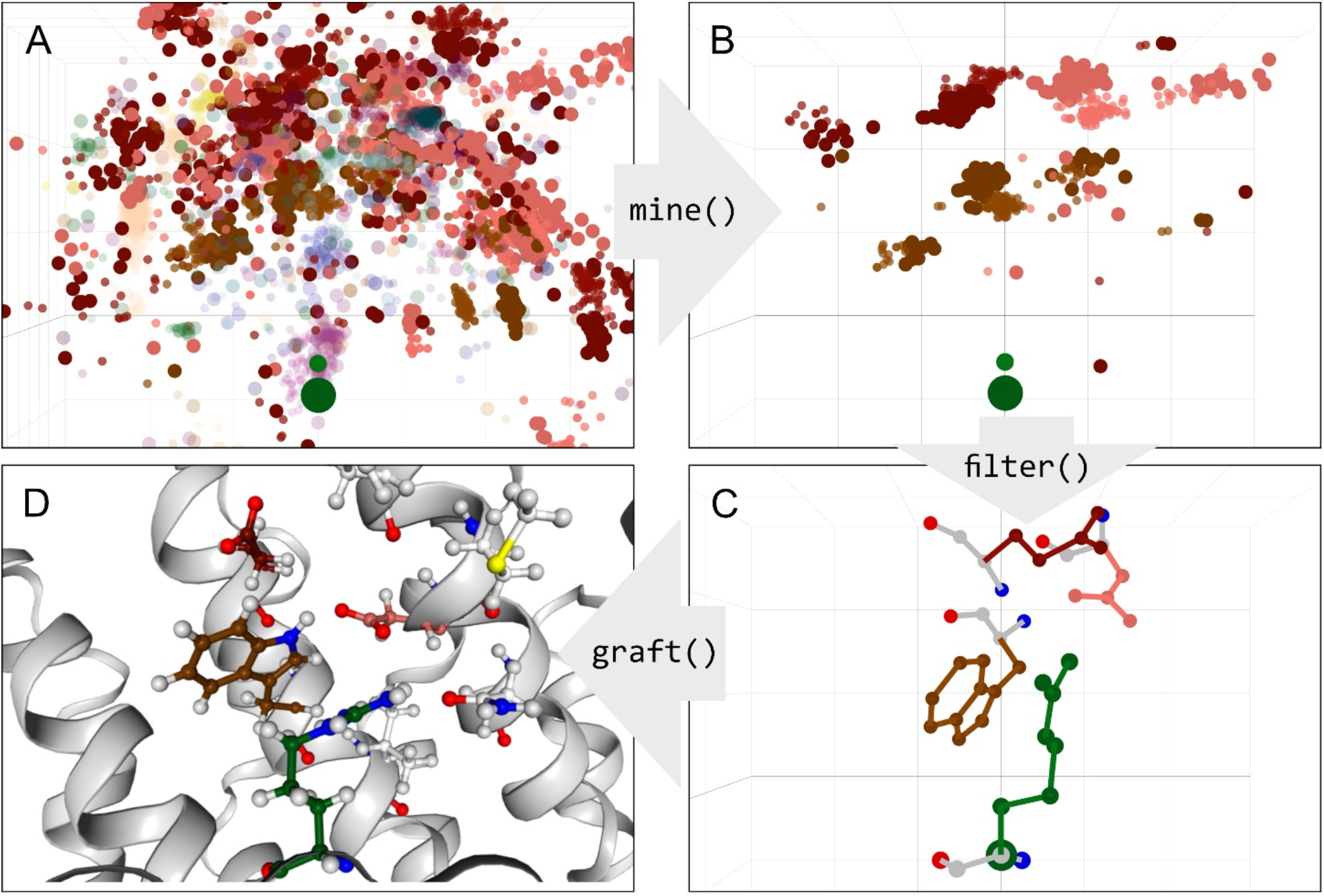
Essential steps during design process with ATLIGATOR. A: 3-dimensional visualization of *Atlas* datapoints for an Arg ligand residue. For reasons of visibility all datapoints with a positive z-value were discarded and only Asp (D), Glu (E) and Trp (W) binder residues are shown with full opacity. B: All datapoints of the DEW *pocket* of an Arg ligand residue derived by mining (A). C: Detailed view on a single instance of the DEW *pocket* including all side chain atoms. D: DEW *pocket* grafted onto an armadillo repeat protein scaffold. The mutated residues are highlighted by coloring their carbon atoms according to the ATLIGATOR scheme. In A, B and C the axes are defined using the standard x, y and z coordinate definitions. The Cα atom of the Arg ligand residue is highlighted by a bigger radius as the center of view. In A and B every binder residue as well as the ligand residue consists of a Cα atom (stronger color, big radius) and a Cβ atom (lighter color, small radius) colored according to the ATLIGATOR color scheme.

##### Pocket Grafting and Quick Graft

Gathered insights and ideas from *atlas* and *pockets* can be applied to a protein of interest to craft designs with new binding features. Pocket grafting and the quick graft protocol can help fulfill this task. By supplying a structure of the protein-protein or proteinpeptide complex of interest as a scaffold and selecting a previously mined collection of pockets such a task can be started. After defining mutable groups of peptide or protein ligand and binder within the scaffold this can be fed into a new design. Here, *pockets* of the assigned *pocket* collection can be selected and grafted automatically onto the binder protein (see 2.1.5 and 2.1.6). If *pockets* are chosen for neighboring ligand residues and the same binder residue is mutated multiple times conflicts may occur. Such conflicts are internally solved based on cumulative similarity scoring (see 2.1.5 and 2.1.6) and provide the optimal grafting solutions. Nevertheless, the mutations are based on natural pockets in the input structures and the side-chain rotamers will not fit perfectly into the new backbone. Thus, we recommend minimizing these rotamers subsequently, e.g. with the Rosetta fixbb side-chain packing protocol (Leaver-Fay *et al*., 2011), to receive a self-consistent representation of all mutable residues. Designs can be written into a pdb file.

## 3 Results and Discussion

There has always been an interest in computational structural biology to describe and classify protein side-chain interactions. The introduction of such descriptions established by Singh & Thornton (Singh and Thornton, 1992) led to improved understanding, but so far the utilization of this data was not focused on protein design approaches. To do so, we created ATLIGATOR to more automatically detect naturally occurring interaction patterns and feed them into the design of protein-protein and protein-peptide interactions.

Now that *atlases* can be generated and designs can be created based on this data, it is interesting to look at the main functions of ATLIGATOR in the context of a typical workflow e.g. when designing a peptide binding interface. As an example, we chose the redesign of the binding interface of a designed armadillo repeat protein (dArmRP) that binds a peptide with the sequence [KR]_5_ (Hansen *et al*., 2016). We will focus on the third arginine in the peptide. On the one hand we aim to improve binding to arginine by pocket grafting and on the other hand we would like to alter the binding preference to isoleucine.

For redesigning the dArmRP binding site we decided to use structures assigned to SCOPe identifier a.118 (alpha-alpha superhelix) as our data source based on their structural similarity to our target protein. We processed all corresponding structures with ALARMS (parameters shown in Table 1). Hereby, we selected 907 structure files from the Protein Data Bank, leading to 2584 processed substructure complexes.

**Table 1:** Parameters for *ALARMS* processing, *atlas* generation and pocket mining.

| ALARMS parameter | Value |
| --- | --- |
| Distance | 8 Å |
| Threshold binder | 40 aa |
| Min ligand | 3 aa |
| Hydrogen interactions | No |
| Atlas generation parameter | Value |
| Min length ligand | 3 aa |
| Max length ligand | 20 aa |
| Interaction radius (default) | 4.0 |
| Interaction radius (h-bond) | 6.0 |
| Interaction radius (aromatic) | 6.0 |
| Interaction radius (Ionic) | 8.0 |
| Skip backbone atoms | True |
| Allow intramolecular interactions | False |
| Pocket mining parameter | Value |
| Max p per Lig Res | 10 aa |
| Minimum Pocket size | 3 aa |
| Confidence Threshold | 0.02 |
| Support Threshold | 0.01 |
| Cardinality Base | 1.21 |
| Distance Factor | 2.0 |
| Orientation Factor | 1.0 |
| Secondary Orientation Factor | 1.0 |
| Variance Threshold | 9999 |

The *atlas* was generated from all structures obtained in the last step using parameters shown in Table 1. The *atlas* includes 20 pages containing 400 maps (see Figure 2**Error! Reference source not found**.) in total - respective to every combination of canonical amino acids – comprising 43752 datapoints. Of these, 4869 datapoints contain Arg as ligand residue (Figure 3A), with the most frequent interaction partners of Arg being Asp (1055), Glu (998), and Thr (482). To get a better understanding of these interactions we analyzed frequent groups of interactions by *pocket* mining using the parameters shown in Table 1. One prevalent motif is the DEW *pocket*, which contains the residues Asp, Glu and Trp (see Figure 3B). An example single *pocket* instance is shown in Figure 3 C. These *pockets* are then used for grafting onto the scaffold of choice as shown exemplarily in Figure 3D for a DEW *pocket* grafted on the dArmRP scaffold. Such designs can now serve as starting points for further calculations or experimental testing.

For our second objective of altering the binding preference to isoleucine, we used the same atlas and pockets as above. Here, 1447 datapoints contained Ile as the ligand residue. The most frequent interaction partners of Ile were Tyr (157), Asp (150), and Met (148). For the transfer of an Ile binding pocket, we want to highlight an Ile – RDFY *pocket*, which was found in 8% of all Ile ligand residues. The RDFY interaction groups that were extracted from proteins that contain the alpha-alpha superhelix fold (a.118) are shown in Figure 4A. The single pocket instance shown in Figure 4B visualizes the interactions of isoleucine with the members of this pocket. Apart from using this pocket for grafting, it can also be used to search for similar binding pockets present in different folds. In fact, when we did this, we found a motif with remarkable similarity in the ankyrin repeat cluster domain 4 of human Tankyrase 2 (Guettler *et al*., 2011) which is unrelated to our input *structure collection*. Interestingly, this motif is interacting with an Ile side chain of a bound peptide in a similar way (see Figure 4C), supporting the idea that general descriptions of binding pockets can exist in different folds. This encourages potential transferability of pockets from one protein to another.

**Figure 4:**
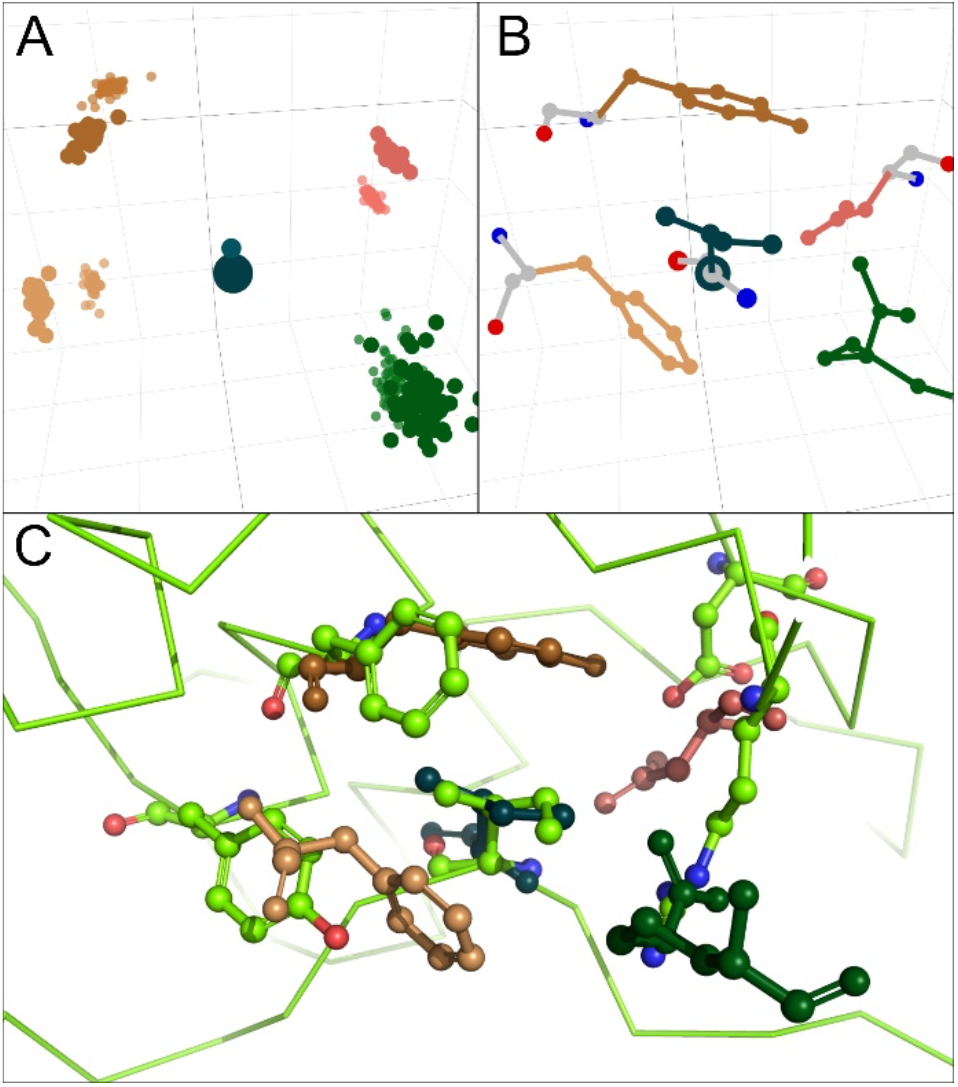
Ile - RDFY *pocket* in comparison to a natural binding pocket found in an unrelated peptide-binding protein. A: Cα and Cβ data points of Ile-binding pocket containing Arg, Asp, Phe and Tyr (see Figure 3B). This *pocket* is based on an *atlas* of structures assigned to SCOPe classification a.118 (alpha-alpha superhelix). B: Single RDFY *pocket* with complete side-chain configuration, originating from pdb structure 2ein. C: Overlay of *pocket* in B (same coloring) and an Ile – RDFY interaction (green) found in an ankyrin repeat protein (d.211.1.0).

Surprisingly, the individual *pocket* instance in Figure 4B does not originate from the alpha-alpha superhelix (a.118) domain, but other parts within the larger multidomain protein. In fact, this binding pocket is formed by three subsections of two polypeptide chains, classified as f.17.2.1, b.6.1.2 and f.23.3.1. This is due to the fact that the original polyprotein complex contains just one a.118 subunit and no additional filtering was applied to input structures for *atlas* generation. Even though the *pocket*’s origin is not the same fold as our scaffold protein, this is a very strong hint that interaction motifs found with ATLIGATOR can be generalized to other folds – even if more than one chain is forming such a binding pocket. Thus, analyzing *atlases* or *pockets* from different origins will help understand relationships of yet uncovered binding motifs.

In fact, ATLIGATOR is a versatile, data-driven methodology to analyze protein-protein and protein-peptide interactions in a variety of protein folds. In contrast to other tools, it focusses on local interactions, basically tracking down the problem onto the side-chain level while incorporating intuitive design options. Moreover, it opens the opportunity to compare binding motifs from different sources to answer questions about generalizability of such motifs. Hence, fold-specific motifs can be detected and compared. ATLIGATOR also features statistical tools which can be utilized for analyzing interactions within the context of an *atlas, atlas map, atlas page* or *pockets*.

Despite these possible applications of ATLIGATOR, the main focus is to analyze the interaction in *atlas* and *pockets* for further use in a specific design task. To this end, it includes multiple ways to visualize and use data stored in the *atlas* and *pockets* and provides pocket grafting and quick graft options enabling a unique use of the interactions leveraged from the input structures.

## Acknowledgements

We thank members of the Höcker lab and the PReART research team for discussions and feedback, in particular Florian Gisdon, Julian Beck, Merve Ayyildiz, Steffen Schmidt, Pascal Kröger, Noelia Ferruz, Dominik Lemm and Abhishek Anan Jalan.

